# Variation in the rate lateral gene transfers accumulate in a grass lineage

**DOI:** 10.1101/2022.10.10.511554

**Authors:** Pauline Raimondeau, Matheus E. Bianconi, Lara Pereira, Christian Parisod, Pascal-Antoine Christin, Luke T. Dunning

## Abstract

Lateral gene transfer (LGT) has been reported in multiple eukaryotes. This process seems particularly widespread in the grass family, although we know very little about the underlying dynamics and how it impacts gene content variation within a species. *Alloteropsis semialata* is a tropical grass, and multiple LGT were detected in a reference genome assembled from an Australian individual. As part of this study we assemble three additional *de novo* genomes for *A. semialata* and one for its sister species *Alloteropsis angusta*. In total we detect 168 LGT across the five reference genomes. Using whole-genome resequencing data for a further 40 individuals we establish the distribution of these transfers and map their origin along the species phylogeny. This shows that many LGTs were acquired relatively recently, with numerous secondary losses. Exponential decay models indicate that the rate of LGT acquisitions varied significantly among lineages [6-28 per Ma], with a high rate of subsequent LGT losses [11-24% per Ma] that largely exceeds that of native loci [0.02-0.8% per Ma]. This high turnover creates large intraspecific structural variants, with a preponderance of LGT occurring as accessory genes in the *Alloteropsis* pangenome. The acquired genes represent unparalleled genetic novelties, having independently evolved for tens of millions of years before they were transferred. Ultimately, the rapid LGT turnover generates standing variation that can fuel local adaptation.

## Introduction

Genomes are dynamic, with continuous gene losses compensated by gene duplications, and occasional *de novo* gene formation (Puigbò et al. 2014; Schlötterer 2015; Murat et al. 2017; Fernández et al. 2020). Because these processes happen at the individual level, they lead to intraspecific variation in gene content. This was originally captured in the concept of the pangenome, developed in prokaryotes which exhibit dramatic gene content variation (Tettelin et al. 2005). In a pangenome, genes are either defined as core, being present in all individuals of a species, or accessory and only present in a subset of individuals. One of the main driving forces for prokaryote pangenome evolution is lateral gene transfer (LGT), which continually adds genetic novelty to a species gene pool (Puigbò et al. 2014; Brockhurst et al. 2019). Pangenomes were more recently established for several groups of eukaryotes, where significant gene content variation was also observed with important ramifications for adaptation (Gordon et al. 2017; Golicz et al. 2020; Tao et al. 2021).

The occurrence of LGT in eukaryotes is now widely accepted (Van Etten and Bhattacharya 2020), and its contribution to the pangenome has been studied in a few groups containing mainly unicellular organisms. Less than 0.5% of the genes in the pangenome of yeast (*Saccharomyces cerevisiae*) have been acquired through LGT (Soanes and Richards 2014), with many of these genes appearing as accessory loci (Han et al. 2021). Similar proportions of LGT have been observed across the phytoplankton (0.16 – 1.44% of the pangenome) and red algae (1%) (Fan et al. 2020), and the distribution of LGT revealed continuous gene transfers during the diversification of the group (Dorrell et al. 2021). LGT also happens in some multicellular eukaryotes (Keeling PJ and Palmer 2008), with unequivocal examples in fungi (Reynolds et al 2018), animals (Xi et al. 2021), and plants (El Baidouri 2014; Li 2014; Li 2018; Wang et al. 2020; Wickell and Li 2020; Ma et al. 2022). Among the latter, LGT is especially prevalent between parasites and their hosts (Yoishida et al. 2010; Kado et al. and Innan 2018; Cai et al. 2021), and in some non-parasitic groups, such as grasses (Mahelka et al. 2017; Dunning et al. 2019; Hibdige et al. 2021; Mahelka et al. 2021; Wu et al. 2022). Some of the LGT received by plants had drastic impacts on the adaptation of the phenotype (Li et al. 2014; Wang et al. 2020; Phansopa et al. 2020), but the temporal dynamics of their LGT and their contribution to the pangenome remain unstudied.

Among grasses, *Alloteropsis semialata* represents an outstanding system to study LGT as a scan of the genome of a single Australian individual identified a total of 59 protein-coding genes laterally acquired from at least nine different donors (Dunning et al. 2019). *Alloteropsis semialata* originated in tropical Africa, where divergent sublineages and the sister subspecies *A. angusta* still occur (Bianconi et al. 2020). Previous analyses have identified LGT shared among multiple individuals of these two species, and even with the more distantly-related *A. cimicina* (Olofsson et al. 2016; Dunning et al. 2019; Olofsson et al. 2019a). However, the reliance on a single reference genome constrained previous systematic detection efforts to LGT present within this individual. Quantifying the rates of LGT turnover and its contribution to the gene content in the species as a whole requires considering multiple reference genomes for a diverse set of accessions, and estimating the ages of LGT based on their distribution among accessions.

In this study, we generate complete reference genomes for three accessions of *A. semialata* representing various African sublineages and one for *A. angusta*, leading to five reference genomes for diploid individuals with the inclusion of the original Australian reference (Dunning et al. 2019). We use phylogenetic approaches to identify all protein-coding genes laterally acquired from other groups of grasses in each of the five genomes. We then use whole-genome sequence data for 40 additional diploid *Alloteropsis* accessions to (i) establish the distribution of all identified LGT across the diversity of the group and map their origin onto a time-calibrated phylogeny, testing for the contributions of shared history and close geographical proximity on the sorting of LGT. The timing of the acquisition of each LGT is then fitted to an exponential decay model to (ii) directly estimate, for each of the five reference genomes, the rate of LGT gain and subsequent loss in their ancestors, and to contrast the latter with the rate of loss of native genes. Finally, (iii) we compare the intraspecific variation in LGT content to the amount of variation of native genes to quantify the contribution of LGT to the pangenome of the group. Overall, our analyses show that high LGT turnovers overly contribute accessory genes to the pangenome.

## Results

### Assembling multiple *Alloteropsis* reference genomes

This study uses five *de novo Alloteropsis* genomes, four of which were sequenced as part of this study. This includes three *Alloteropsis semialata* assemblies that, together with the previously sequenced genome from an Australian individual (accession AUS1; Clade IV; Dunning et al. 2019), encompass the four main nuclear clades within this species (Bianconi et al. 2020): one individual from South Africa (accession RSA5-3; Clade I), one from Tanzania (accession TAN1-04B; clade II), and one from Zambia (accession ZAM1505-10; clade III). All four *A. semialata* accessions were sequenced with a hybrid approach combining short-read Illumina and long-read PacBio data. We also generated an assembly for *Alloteropsis angusta*, the sister species of *A. semialata*. For the *A. angusta* genome we used an individual from Uganda (accession AANG_UGA4), and this was the only accession solely sequenced with short-read Illumina data (Table S1).

The size of the *A. semialata* assemblies ranged between 0.62 and 0.86 Gb, which reflects differences in the genome size estimates based on flow cytometry for these diploid accessions (Bianconi et al. 2020), although the mean assembly size was 27% (range 20 - 32%) smaller than the flow cytometry estimates (Table S2). Accessions RSA5, TAN1 and AUS1 (prior to super-scaffolding using Dovetail Genomics Chicago and Hi-C data) had similar assembly statistics: mean N50 = 0.18 Mb (SD = 0.01 Mb); mean number of scaffolds = 7,059 (SD = 1,018), mean longest scaffold = 1.07 Mb (SD = 0.08 Mb; Table S2). The ZAM1505-10 accession was generally more fragmented than the other three *A. semialata* reference genomes: N50 = 0.07 Mb; number of scaffolds = 19,813; longest scaffold = 0.60 Mb (Table S2). BUSCO analyses indicated that all four *A. semialata* genomes were comparably complete (mean = 89.3%; SD = 2.3%), although the ZAM1505-10 assembly had higher levels of duplication (25.5%) compared to the other three (mean = 7.3%; SD = 1.8%; Table S2). This same pattern was repeated when the BUSCO analysis was performed solely on the genome annotations, with a comparable number of complete annotations among assemblies (mean = 84.7%; SD = 3.5%), and with ZAM1505-10 having a higher level of duplicated annotations (26.3%) when compared to the other three *A. semialata* (mean = 7.5%; SD = 1.7%; Table S2). The increased annotation duplication in ZAM1505-10 is accounted for by our downstream LGT identification pipeline.

The AANG_UGA4 *A. angusta* assembly was more fragmented and less complete than the four *A. semialata* references, largely owing to the lack of long-read PacBio data for this accession. The initial Illumina assembly was only marginally smaller (0.92 Gb) than the flow cytometry estimate (0.97 Gb; Bianconi et al. 2020), but the N50 (20 kb) and number of scaffolds (179,640) indicated it was highly fragmented (Table S2). We therefore re-scaffolded the AANG_UGA4 assembly in relation to the chromosomes of the AUS1 accession prior to genome annotation. According to the BUSCO analysis, the AANG_UGA4 assembly was only 77.1% complete, and this value decreased to 67.4% when considering the genome annotations. Despite the lower quality of this assembly, it is unlikely to bias our results as it effectively acts as an outgroup to the *A. semialata* accessions.

### Widespread lateral gene transfer in *Alloteropsis*

For each of the five reference genomes we independently detected all protein-coding LGT using an approach we previously developed (Dunning et al. 2019; Hibdige et al. 2021), which identifies genes nested in distantly-related phylogenetic groups before applying successive filters to verify that a scenario of LGT is statistically supported (Figure 2). As the method was applied to each genome independently, older shared LGT would potentially be detected in multiple scans. We therefore generated a merged list ensuring the occurrence of each LGT was only counted once. Our method also counts recent duplicates (i.e defined as being part of the same monophyletic LGT cluster in the phylogeny) as a single LGT. This negates the problem of having to determine whether the gene duplicate arose prior or post lateral transfer, or indeed whether the duplication is in fact an assembly error. In total, we identified an initial 177 LGT (both primary and secondary candidates *sensu* Dunning et al. 2019) across the five Alloteropsis genomes (Table S3; see Supplementary Results).

Genome assemblies and annotations are seldom complete, and there will often be missing loci. To reduce the effect of assembly and annotation quality on our results we decided to verify if each LGT was present in the unassembled sequencing reads using a phylogenetic approach. For 11 LGT, individual reads could not be reliably assigned to specific gene copy, and these LGT were not considered further in this study. For two of the remaining LGT, in depth phylogenetic analyses indicated that the genes had been transferred twice independently to different accessions of *A. semialata* (Figure S6 & S7; see Supplementary Results), and the genes resulting from each of these events were considered as independent LGT, leading to a final total of 168 LGT (range of 32 – 100 per genome; Table S3). These LGT were acquired from three different subfamilies: 92.2% Panicoideae, 4.8% Chloridoideae, and 3.0% Danthonioideae (Figure 2; Table S4). Within the Panicoideae the main donors were Cenchrinae (n = 88 LGT), Andropogoneae (n = 54), and Melinidinae (n = 7). In numerous cases the LGT form clusters in the assembled genomes of *Alloteropsis* (Figure S1; see Supplementary Results), confirming that multigene genomic blocks can move between different groups of grasses (Dunning et al. 2019).

### History and geography both explain the distribution of LGT among accessions

Using a sequencing read analysis, the distribution of each of the 168 LGT was established among 40 additional accessions of *Alloteropsis*, including numerous *A. semialata* and *A. angusta* individuals, as well as the outgroup *A. cimicina* (Figure 1). Together with the five reference genomes, these 45 accessions cover the known geographical and genetic diversity of the genus, with divergence times spanning more than 11 million years (Figures 1 and 2). Only two LGT were detected in 44 accessions (Figure 2a) and were likely acquired before the diversification of *Alloteropsis*, with few other LGT shared across species (Figure 1). A majority of the LGT were restricted to phylogenetic subgroups, with five LGT only present in a single individual (ZAM1505-10; Figure 1; Table S4), suggesting very recent acquisitions. The number of LGT shared among pairs of individuals decreased with divergence time (Mantel test, p-value < 0.0001, R^2^ = 0.53; Figure S2), which supports a gradual accumulation of LGT during the diversification of *Alloteropsis*. We conclude that history largely explains the patterns of LGT distribution within and among *Alloteropsis* species.

**Figure 1:**
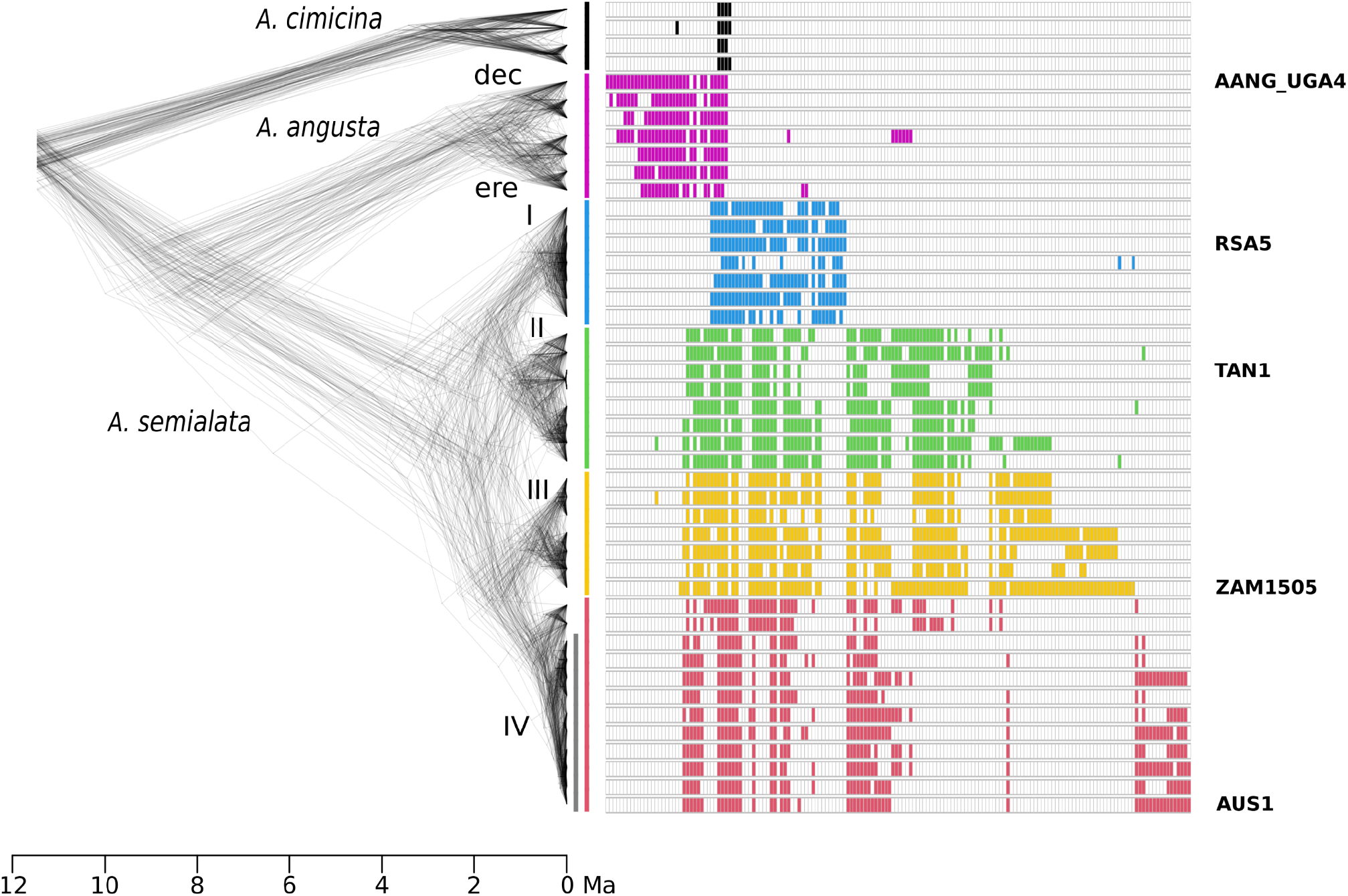
Phylogenetic distribution of laterally acquired genes. Phylogenetic relationships are shown on the left, with each time-calibrated phylogenetic tree based on a different set of five nuclear genes. The phylogeny tips are ordered as in Table S4. The time scale, in million years, is shown at the bottom. The species names are indicated near branches corresponding to their most recent common ancestor, as are those of the groups corresponding to the two ecotypes of *A. angusta* (‘ere’ = erect, ‘dec’ = decumbent [Curran et al. 2022]), and the subclades of *A. semialata* (clades I to IV [Bianconi et al. 2020]). Coloured bars on the right of the phylogenetic trees delimit the clades, with the grey bar within clade IV highlighting the accessions from Asia and Australia, when all others originate from Africa. The presence of each LGT in an accession is indicated with a rectangle coloured according to the clade; black = *A. cimicina*, purple= *A. angusta*, blue = clade I of *A. semialata*, green = clade II of *A. semialata*, orange = clade III of *A. semialata*, red = clade IV of *A. semialata*. The LGT are ordered by their abundance in each group as in Figure S1, which highlights physically-linked LGT, and Table S4, which gives detailed information for each LGT. The names of the reference genomes are shown on the right.

**Figure 2:**
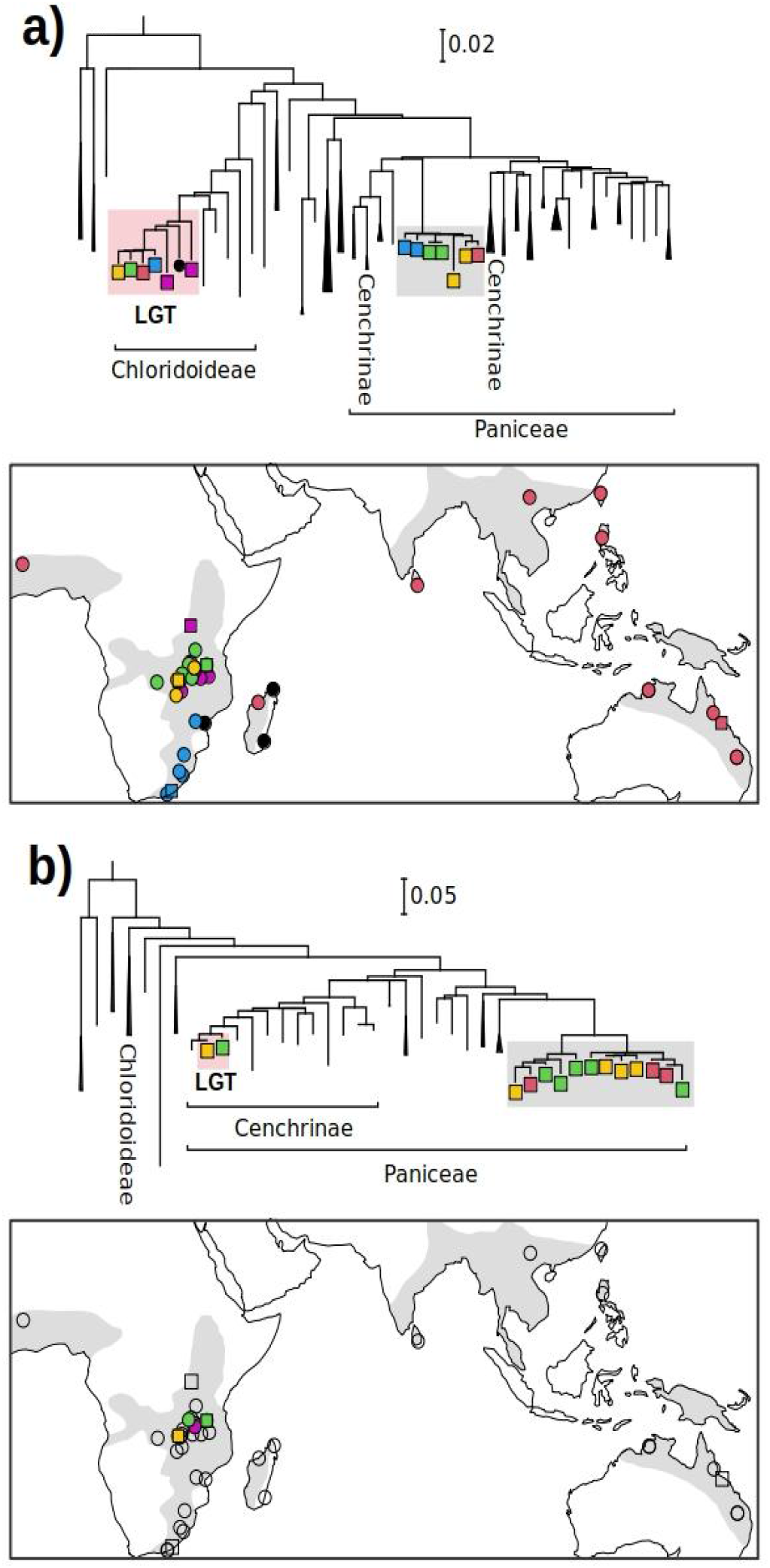
Phylogenetic evidence and geographical distribution of two exemplar laterally-acquired genes. These two genes were selected to represent **a)** a lateral gene transfer (LGT) present in most individuals (LGT-001, reference gene ASEM_ZAM1505-10_31837) and **b)** one restricted to a few individuals (LGT-103, reference gene ASEM_TAN1_34267). For each of them, a simplified phylogenetic tree at the top shows the positions of the laterally acquired genes of *Alloteropsis* (“LGT”; highlighted with pink shading) and the native copies (highlighted with grey shading). In both groups, the position of the genes extracted from *A. cimicina* and the five reference genomes of *A. semialata* and *A. angusta* is indicated with colours matching Figure 1. The donor clades are delimited at the bottom. For each LGT, the geographical position of accessions bearing them are shown on maps with filled squares (for reference genomes) or filled circles (for resequenced genomes), coloured by phylogenetic groups as in Figure 1. The positions of individuals without the LGT are shown with empty squares (for reference genomes) and empty circles (for resequenced individuals). The grey shading represents the known distribution of *A. semialata*.

Although evolutionary history appears to be the main driver of present-day LGT distribution, geography also appears to play a role. Some LGT present a patchy distribution across the *Alloteropsis* phylogeny, being present in a few accessions belonging to different phylogenetic groups (Figure 1). These patterns could result from repeated losses following ancient acquisitions, but in some cases likely result from introgression of recently-acquired genes. Indeed, some of these LGT with a patchy distribution are shared among distantly related accessions, sometimes from different *Alloteropsis* species, which cluster geographically (Figure 2b; see Supplementary Results). Once divergences times were accounted for, the number of LGT shared by pairs of accessions decreased with geographical distances (partial Mantel test, p-value = 0.0006, R^2^ = 0.08; Figure S2), showing that history and geography both contributed to the present-day distribution of LGT.

### The rate of LGT gains varies among lineages

The origin and potential loss of each of the 168 LGT were individually inferred along one of a hundred time-calibrated phylogenetic trees that best explained the LGT history, using stochastic mapping with 100 pseudoreplicates. The origin of each LGT was then retrieved for the line of ancestors leading to each of the five reference genomes containing it. When more than one origin of the LGT was inferred by the stochastic mapping, the LGT was discarded (ranging from 25% to 56% of LGT; Table S3). The inference of multiple origins might result from repeated losses of ancient LGT or spread of recent LGT via introgression, but in both cases, the patchy distribution of the LGT across the phylogeny means that the LGT event cannot be confidently assigned to a single time point.

As expected given the random nature of the stochastic mapping approach, the inferred ages of origin varied among the 100 replicates, but there was a consistent increase of observed LGT origins toward the present leading to the ZAM1505-10 genome, and to a lesser extent to some of the others (Figure S3). When considering the average across pseudoreplicates, the number of observed LGT increased strongly toward the present in RSA5, AUS1, and especially ZAM1505-10, where it fitted closely to an exponential distribution (Figure 3a). Based on fitted exponential decay models, the rates of gains were estimated between 2.66 and 15.74 LGT per Ma for the five genomes. The highest value was observed in ZAM1505-10, with a rate of gains significantly higher than the others, which all had overlapping confidence intervals for this parameter (Figure 3b). If the LGT for which ages could not be reliably estimated follow the same distribution as the others, the rate of gains would be inflated to between 3.54 and 28.1 LGT per Ma (in ZAM1505-10; Figure 3b).

**Figure 3:**
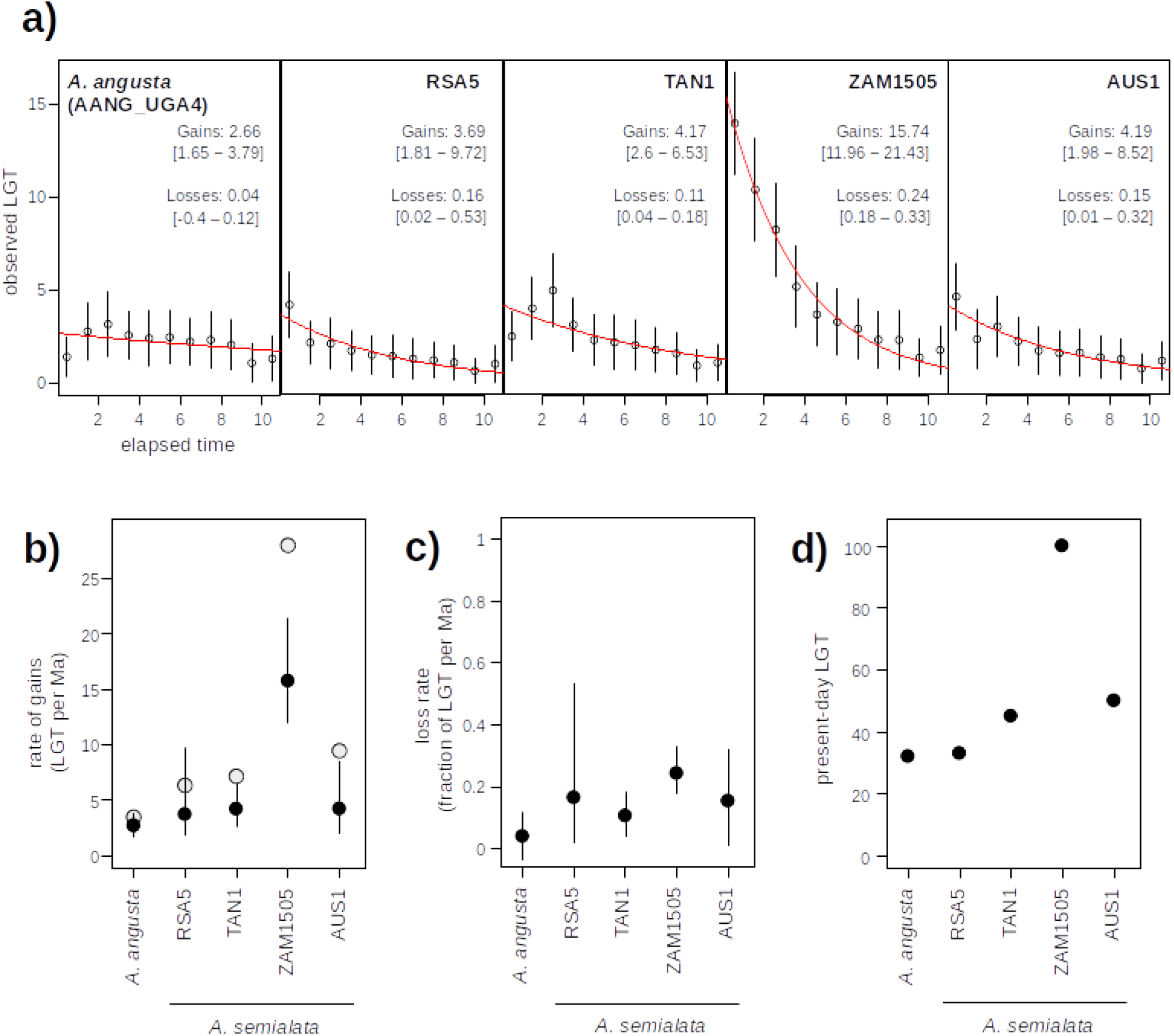
Rates of gains and losses of lateral gene transfers. **a)** Numbers of lateral gene transfers (LGT) assigned to different 1 Ma time slices. For each of the five reference genomes, named at the top, the points indicate the mean and the bars the 95% intervals across 100 pseudoreplicates. The red curves show the fitted exponential decay model. The inferred rates of LGT gains and losses per Ma are indicated, with 95% confidence intervals in square brackets. **b)** The inferred rate of LGT gain per Ma is shown with filled circles for the five references genomes, with segments showing the 95% intervals across replicates. The opened circles show the estimates obtained assuming that LGT for which a time of origin could not be estimated are distributed through time proportionally to those with estimated ages. **c)** The inferred rate of LGT losses per Ma is indicated for the five reference genomes. The points show the mean and the bars the 95% intervals across 100 pseudoreplicates. **d)** The number of LGT observed in the genome of each of the five reference genomes is indicated.

The rate of loss of LGT per Ma in *A. angusta* was estimated to be 4%, mirroring the weak increase of LGT toward the present (Figure 3a). The modelled relationships indicated that between 11% and 16% of LGT were lost each Ma in the lineages leading to most *A. semialata*, while 24% were lost in the lineage leading to ZAM1505-10, although the confidence intervals of the latter overlap with those of some others (Figure 3c). The high number of LGT present in the genome of ZAM1505-10 therefore results from a higher rate of gains, despite a potentially higher rate of losses (Figure 3). Overall, these analyses indicate that the genomes of *Alloteropsis* undergo a high turnover of LGT, with repeated gains throughout their history followed by relatively rapid losses. The rates of gains and losses however vary among sublineages, creating important variation in LGT content, especially within *A. semialata*.

### LGT are lost faster than native genes

To compare the rate of LGT loss to that of non-LGT components of the genome, we used a dataset of 6,657 relatively conserved genes that were present as a single copy in the common ancestor of *Alloteropsis* and other Paniceae, and that did not have a signal of being LGT. Using the same sequencing read analyses as for the LGT, we estimated the number of native genes lost every Ma, for the lineages leading to each of the five reference genomes. Assuming that the losses whose age could not be estimated are proportionally distributed among the time windows, the fraction of the 6,657 genes lost every Ma ranges from 0.25% in *A. angusta* to 0.04-0.06% in the four *A. semialata* (see Supplementary Results). We conclude that LGT are lost up to 500 times faster than native components of the genome of *A. semialata*.

### LGT overly contribute to gene presence / absence variation

We evaluated the pangenome variation focusing on the five reference genomes, as genes specific to other accessions cannot be identified based on the data available. Out of 168 LGT, only two are present in all five *A. semialata* and *A. angusta* reference genomes, and can thus be considered as core genes, with the vast majority (166 genes; 98.8%) appearing as accessory genes. Of the 166 polymorphic LGT, 21 are specific to *A. angusta*, while 136 are specific to *A. semialata*, with the other 9 shared by some accessions of the two groups (Figure 4). Of the LGT specific to *A. semialata*, 130 vary within *A. semialata*, and most of the pangenome variation was generated by LGT restricted to ZAM1505 and to a lesser extent AUS1 (Figure 4). These results show that recent LGT gains and losses create important pangenome variation, within *A. semialata* and among the two sister species. In contrast to LGT, only 3.4% of the 6,657 native genes investigated above appear as accessory, and this is mainly due to numerous losses in the *A. angusta* lineages, which contributes in excess to the native pangenome (Figure 4). This result is not due to the more fragmented and incomplete nature of the *A. angusta* reference genome as this analysis was performed on the unassembled reads. Within *A. semialata*, the four accessions contribute in similar proportions to the native pangenome variation, which stands in stark contrast to the LGT pangenome contributions (Figure 4).

**Figure 4:**
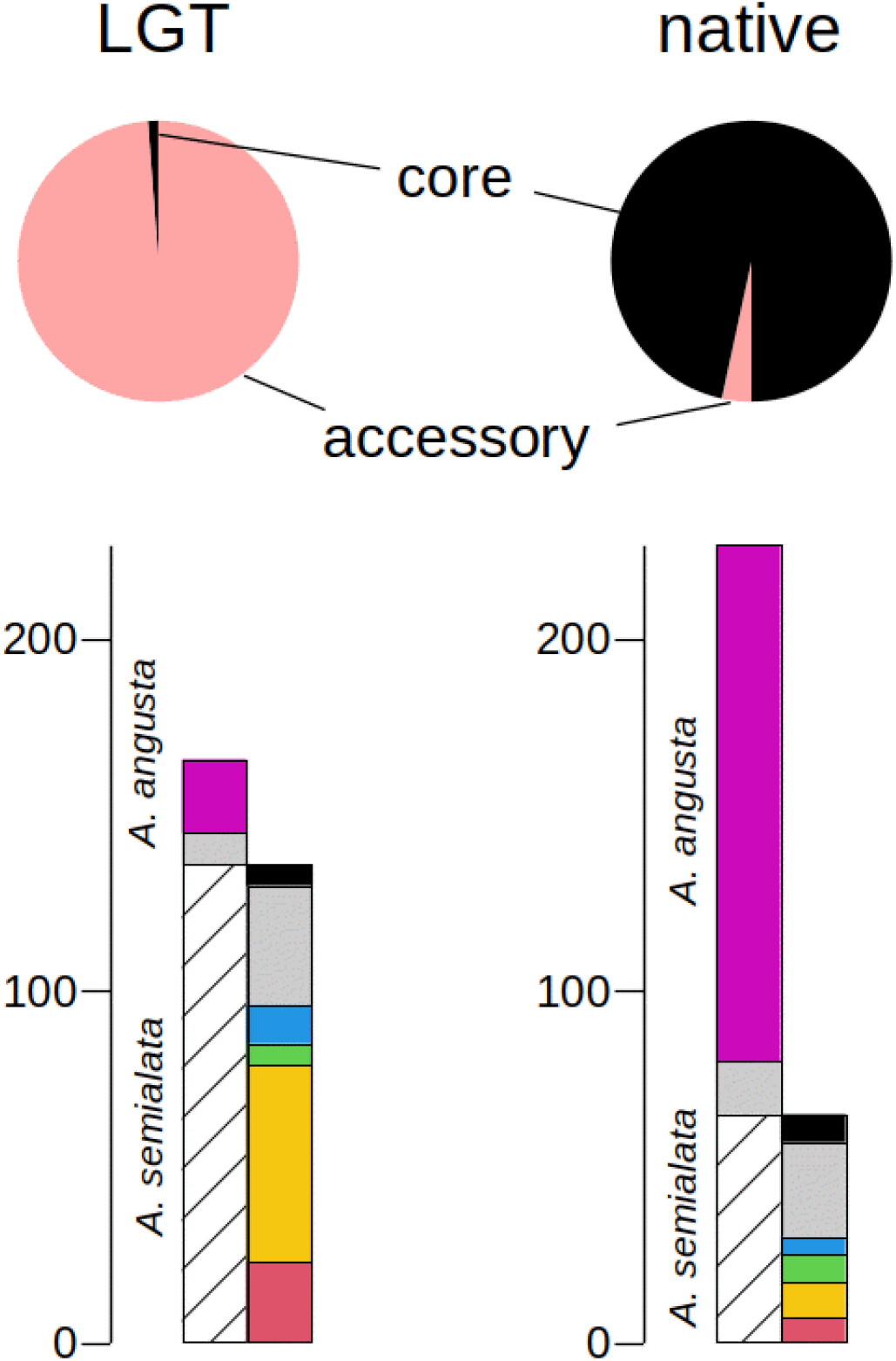
Contributions of laterally acquired genes to the pangenome of *Alloteropsis*. Pie charts at the top show the proportion of accessory genes (variable among the five reference genomes), for genes resulting from lateral gene transfers (LGT) and for native genes inherited vertically from the *Alloteropsis* ancestors shared with other Paniceae grasses. For both LGT and native genes, the contributions of each group to the pangenome variation are indicated, with the number of variants specific to *A. angusta* indicated with purple bars, the number specific to *A. semialata* in black hatching, and the number of variants occurring in both species in grey. For *A. semialata*, the source of accessory genes is further decomposed, with those fixed within the species in black, those variable and shared among multiple accessions in grey, and those restricted to one accession with bars coloured as in Figure 1; blue = RSA5, green = TAN1, orange = ZAM1505, red = AUS1.

Our analysis of native genes was conservative, and we consequently captured as few as one tenth of all annotated protein-coding genes (Table S1). If the proportion of variable genes was maintained, we would thus expect up to 2,270 accessory native genes in the pangenome represented by the five reference genomes analysed here. This is a lot lower than the number of accessory genes identified in other grass pangenomes, such as maize where 69% of 103,033 pan-genes identified from 26 reference genomes had a variable distribution (Hufford et al. 2021). However, a majority of these variable genes in maize only share orthologs with close relatives or are in fact maize specific (Hufford et al. 2021), meaning they would not be included in our conservative phylogenetic approach. It is also worth noting that our analyses are based on read presence/absence and not gene annotation presence/absence. This means that our approach is again conservative as it is not as affected by genome assembly and annotation quality, and it also counts pseudogenes still present in the genome. Whilst these methodological and data volume differences (>5x fewer *Alloteropsis* than maize assemblies) result in a fewer number of variable pan-genes in *Alloteropsis* compared to other species, our pipelines to trace the evolutionary history of LGTs and native genes is equally conservative and is therefore comparable. In total, 99% of detected LGT in *Alloteropsis* are variable, which equates to 166 LGT in total. The number of LGT is also likely to be underestimated, but they still represent at least 7% of the variable pan-genes, despite accounting for less than 0.5% of all protein-coding genes. Overall, these patterns indicate that, despite their low relative numbers, the LGT disproportionately contributes to the pangenome of *Alloteropsis*, and that this excess contribution stems from both a high rate of LGT gain and secondary loss.

### Function of the LGTs

The final list of 168 LGTs were annotated against the SwissProt database revealing a diverse set of functions, including genes associated with C_4_ photosynthesis, disease resistance and abiotic stress tolerance (Table S5). A subsequent GO enrichment analysis identified three significantly (adjusted p value < 0.05) overrepresented categories: cellulose biosynthetic process, cellulose synthase (UDP-forming) activity, pre-mRNA 3’-splice site binding and ribonuclease P activity (Table S6). It is unsurprising that there are relatively few enriched categories when all LGT are considered, as they include both the genes that were potentially selected for in addition to adjacent loci that hitchhiked along as part of the same fragment of DNA (Olofsson et al. 2019a). Further work to disentangle the two may provide clarity as to whether genes with certain functions are preferentially retained after the initial transfer.

## Discussion

### Continual birth and death of lateral gene transfers

We can only detect laterally-acquired genes (LGTs) that are retained until the present. The detected LGTs therefore likely represent the tip of the iceberg of a much higher background rate of transferred genes that were then lost through drift or negative selection. Comparison of LGT content among multiple accessions of the same group can help infer the dynamics of gene transfers through time. For instance, LGT restricted to accessions of *Alloteropsis semialata* from Australia presenting a high similarity with populations of the donor species from the same region must have been acquired after the colonization of Australia, inferred in the last 0.5 Ma (Olofsson et al. 2019). Conversely, those LGT present in all *Alloteropsis* species which accumulated substitutions after the transfer (Figure 2a) were likely acquired before the diversification of the genus some 11 Ma. The exact origin of a majority of LGTs is difficult to precisely date, but their distribution among accessions can be used to model the temporal spread of acquisitions, but also of subsequent losses.

The LGT present in a given genome represent those that were gained and persisted minus the proportion that were lost over time. Assuming that the rate of gain, corresponding to the rate of integration of foreign gene in the genome of *A. semialata*, and the rate of subsequent losses are constant through time, the number of LGT persisting at present should decrease with their age, following an exponential decay model. Such patterns were indeed observed for the LGT found in the genomes of *A. semialata* accessions (Figure 3a). These analyses bring direct support for the accumulation of LGT in the genomes of plants, and also indicate that between 6 and 28 LGT entered the genomic lineages of the four *A. semialata* analysed here every million years (Figure 3b). However, between 11% and 24% of those LGT originally integrated are lost every million years, a proportion that vastly exceeds the fraction of native genes lost every million years (0.04-0.06% for *A. semialata*; Figure S4). These numbers indicate a high LGT turnover rate, with potentially more than 20 foreign genes entering the recipient gene pool every million years, but half of those LGT being lost after 3-6 Ma.

### Lineage specific dynamics of LGT gains and losses

While our models assume constant rates of LGT gains and losses through time, this assumption is likely violated. First, the raw numbers of DNA transfers are likely to vary as transfer opportunities fluctuate. Such variation might represent changes in the phenotype of the recipient. For instance, high levels of selfing would be expected to reduce LGT via illegitimate pollination, whereas increased vegetative propagation would provide more opportunities for LGT through root inosculation. In addition, the probability of DNA transfers likely depends on the presence of potential donors in close contact, which can vary as species migrate. Second, the probability that transferred DNA persists over generations depends on the demography of the recipient species. Indeed, large effective population sizes would increase the chance of advantageous LGT spreading under selection, while increased drift in small populations would increase the retention of neutral LGT. The causal factors are difficult to identify, but such variation is likely responsible for both deviations from the models within each lineage and differences among them.

The most striking difference concerns the lineage of ZAM1505, which accumulated significantly more LGT than the other lineages (Figure 3). This accession comes from a region of Zambia where *A. semialata* was often found forming multispecies clumps with some known donors (Figure S5), potentially providing increased opportunities for LGT. In addition, this accession is located near the inferred centre of origin of the species (Bianconi et al. 2020), and constant and large population sizes might have favoured the integration of beneficial LGT, as well as physically linked neutral LGT as reported in Australia (Olofsson et al. 2019). These scenarios remain speculative, but the patterns reported here show that some lineages overly contribute to the LGT content of a species.

The precise transfer mechanism is currently unknown, although it has been shown that there is a slight increase in the number of LGT detected in species that produce rhizomes (Hibdige et al. 2021). This could mean that LGT are passed as a result of root-to-root inosculation, as has previously been hypothesised (Dunning et al. 2019). However, LGT is not restricted to grasses that possess rhizomes (Hibdige et al. 2021), meaning that a more general mechanism might underlie the frequently observed transfers. Grasses are wind pollinated, and it has also been proposed that transfers could be a result of reproductive contamination through illegitimate pollination (Christin et al. 2012). This would potentially mirror plant transformation techniques such as repeated pollination (Shan et al. 2005) or pollen tube pathway-mediated transformation (Ali et al. 2015) where the reproductive process is effectively contaminated with DNA from a third individual. These transformation methods require minimal human intervention and could therefore occur naturally, driving the observed grass-to-grass LGT.

### LGT excessively contribute to pangenome variation

We show here that LGT, which represent less than 1% of all genes present in a given accession of *Alloteropsis*, excessively contribute to both the pangenome of *A. semialata* and the joint pangenome of *A. semialata* and *A. angusta* (Figure 4). Because they are continuously acquired and then lost more rapidly than native genes (Figures 3 and S4), most LGT are indeed variable, within the species and even within some populations (Figure S1) (Olofsson et al. 2016). Our results moreover show that even if LGT are preferentially acquired by a given sublineage, such as ZAM1505, they can greatly contribute to the species-level pangenome (Figure 4) and might later be introgressed to other populations (Figure 1; see also Olofsson et al. 2016). Indeed, the different sublineages of *A. semialata* and even the two sister species *A. angusta* and *A. cimicina* occasionally undergo gene flow (Olofsson et al. 2016; Curran et al. 2022), so that the standing variation created by LGT has the potential to fuel adaptation throughout the group.

Unlike duplicates of native genes, LGT have diverged from the other genes in the genome for at least as long as the divergence time between the donor and the recipient, which in the case of LGT reported here extends to more than 40 Ma (Christin et al. 2008). These LGT consequently add diversity to the recipient genomes, both in terms of expression patterns (Dunning et al. 2019) and coding sequences affecting the catalytic properties of the encoded enzymes (Phansopa et al. 2020). We therefore conclude that the high LGT turnover revealed here creates important pangenome variation, and therefore impacts the evolutionary potential of species undergoing such DNA exchanges.

## Conclusions

Grasses appear to frequently undergo lateral gene transfer. Here, we detect LGT from five *de novo* reference genomes belonging to genetically divergent sublineages within the grass *Alloteropsis semialata* and its sister species *A. angusta*. We identify a total of 168 laterally acquired genes, but only two are shared by all five genomes, and the distribution of LGT among 45 *Alloteropsis* individuals suggests a few old acquisitions and many recent ones that are restricted to sublineages. Analyses of the LGT distribution among individuals in a phylogenetic context allowed estimates of the rates of LGT gains and subsequent losses, using an exponential decay model. We estimated that up to 28 LGT per Ma were accumulated by one lineage of *A. semialata*, with up to one quarter of them subsequently lost every million years. The rate of LGT accumulation varied drastically among the five accessions, potentially reflecting differences in LGT opportunities. This high LGT turnover created important inter- and intraspecific variation in LGT content, with almost all LGT being polymorphic, compared to only 3.4% of native genes. LGT therefore excessively contributes to the pangenome variation in this group. Because these LGT provide novelty to the recipient genomes and can be subsequently introgressed among related species, the standing variation revealed here has the potential to fuel rapid adaptation in these grasses.

## Materials and Methods

### Genome sequencing, assembly and annotation

DNA was extracted from live plants grown at The University of Sheffield using the DNeasy Maxi Kit (Qiagen). Short-read Illumina library preparation and sequencing (HiSeq 2500 or 3000) was undertaken at the Edinburgh Genomics Centre (see Table S1 for per sample sequencing details). Long-read PacBio sequencing was generated using the SMRT PacBio platform at the Centre for Genomic Research at the University of Liverpool (see Table S1 for per sample sequencing details). Raw sequencing data were cleaned, assembled and annotated using the same approach as Dunning et al. (2019), full details provided in the supplementary methods.

### Identification of laterally acquired genes

To identify LGTs in each of the reference genomes, we used the phylogenetic method developed in Dunning et al. (2019) and Hibdige et al. (2021), with a few modifications. In brief, this method initially identifies primary LGT candidates as those nested in distantly related groups of grasses in phylogenies of increasing species representation. Secondary LGT candidates are then identified as genes in close physical proximity to the primary candidates that have gene tree topologies supporting the same LGT scenario. The identification of secondary LGT effectively rescues loci discarded by the initial stringent LGT scan. All subsequent analyses made no distinction between primary and secondary LGT. Full details of the LGT identification methods are provided in the supplementary methods

### Distribution of LGTs across *Alloteropsis*

After collapsing recent duplicates (i.e. monophyletic LGT cluster in the phylogeny) we selected a single gene to act as a representative of that LGT (n = 177). The representative was selected based on the original gene tree alignments, with a preference for genes that yielded alignments with the most *Alloteropsis* LGTs present, had the most taxa, and were the longest. In all but two cases a single reference LGT was sufficient to represent the entire LGT group, with two non-overlapping annotated sequences used as references where this was not possible. We then determined the presence of each LGT representative in whole genome short-read datasets belonging to 45 diploid *Alloteropsis* accessions (including the five individuals with reference genomes; Bianconi et al. 2020), using a combination of Blastn searches (minimum alignment length 100 bp) and phylogenetic analyses. For each of the 45 datasets, putative reads corresponding to each LGT were identified via a Blastn analysis with default parameters. The ten top hits were retrieved, and each one was successively aligned with the reference sequences using MAFFT v.7.427 (Katoh and Standley 2013) with the ‘add fragments’ parameter, and a phylogenetic tree was inferred with phyml, with the best substitution model identified using Smart Model Selection SMS v.1.8.1 (Lefort et al. 2017). Reads were considered as belonging to the LGT if they were sister to the reference LGT in this phylogenetic tree. The LGT was considered as present if it was supported by at least three such reads. For the two LGT with two non-overlapping references, the gene was considered as present if either of the reference fragments fulfilled the criteria.

### Molecular dating *Alloteropsi*s

We generated 100 different dated phylogenetic trees, each inferred from 5 randomly selected Benchmarking Universal Single-Copy Orthologs (BUSCO) that could be used to retrace the evolutionary history of each LGT. Consensus sequences for each BUSCO were generated using a reference-based approach (Olofsson et al. 2016; Dunning et al. 2019; Olofsson et al. 2019b), with the chromosome-scale AUS1 *A. semialata* genome as the reference (Dunninget al. 2019). First, we identified BUSCOs from the poales_odb10 database (n=4,896 genes from 12 Poales species) present in the AUS1 genome using Blastp v.2.8.1+. We only considered BUSCO genes that had the same top-hit AUS1 gene for each of the representative species in the poales_odb10 database (n=4,039). The short-read data for each *A. semialata* sample were then mapped to the AUS1 reference using Bowtie2 v.2.3.4.3 (Langmead Salzberg, 2012) with default parameters, except that the maximum insert size for pair-end reads was increased to twice the insert size if necessary. Consensus sequences for each of the BUSCO coding sequences were then called using the methods described in Dunning et al. (2020), with depth filters for each position and individual base calls depending on the coverage of the short-read dataset used (minimum site coverage / minimum base coverage: genome skimming [∼1x coverage] = 1/1; resequencing [∼10x cov.] = 3/2; high-coverage [∼50x cov.] = 10/3). Each of the individual BUSCO alignments was then cleaned using trimAl v.1.2rev59 (Capella-Gutiérrez et al. 2009) with the -automated1 option, which optimises alignments for maximum-likelihood phylogenetic tree reconstruction. We then removed short sequences (< 200 bp) before discarding entire gene alignments that were < 500 bp or did not include all samples. A total of 1,077 gene alignments were then retained.

Time-calibrated species phylogenetic trees were inferred under a coalescence model, using Bayesian inference as implemented in *BEAST2 v. 2.6.4 (Bouckaert et al. 2019). To produce a set of trees representative of the various histories existing within the genomes, analyses were performed on 100 sets of five randomly chosen BUSCO genes. The DNA substitution model was set to a GTR+G, which is reliable across most genes (Abadi et al. 2019), and a relaxed molecular clock with a log-normal distribution of rates was used. To obtain a species tree indicating the relationships among accessions, each individual was set as a different species. For the multispecies coalescent, a constant population function prior was used, and a Yule model was used for the species tree prior. The monophyly of the outgroup (*A. cimicina*) and the ingroup (*A. semialata* plus

*A. angusta*) was enforced to root the tree. The root of the tree was set to 11.46 Ma, using a normal distribution with a standard deviation of 0.0001. This date corresponds to the age estimated based on plastomes (Lundgren et al. 2015) and matches the divergence time of most nuclear genes (Dunning et al. 2017). For each set of five genes, two independent analyses were run for 400,000,000 generations, sampling a species tree every 50,000. After setting the burn-in period to 100,000,000 generations, we used the *tracerer* R package v. 2.2.2 (Bilderbeek and Etienne 2018) to verify that the estimate of each parameter fell within the 95% confidence interval of the other run, indicating convergence. The same package was used to estimate effective sample sizes (ESS) for all parameters based on post-burnin trees combined across the two runs, and 200,000,000 generations were added to analyses where the ESS was below 100 for at least one parameter, until all parameters had ESS > 100. Three analyses did not reach satisfactory ESS after 18,000,000,000 generations, and these were replaced with three other sets of five genes. For two other analyses, the burn-in period was manually adjusted after inspecting the traces. For each set of five genes, the posterior trees from the two runs sampled after the burn-in period were pooled and a summary tree was produced, mapping the median ages on nodes of the highest credibility tree. These trees were visually compared using densitree (Bouckaert 2010).

### Tests for the effects of history and geography on the distribution of LGT

For each pair of individuals, an LGT similarity index was computed as the number of shared LGT. These similarities were statistically compared first to the pairwise divergence times, using a Mantel test. The residuals of this relationship, which represent the part of the shared LGT not explained by divergence times, was then tested for an effect of pairwise geographical distance, using the partial Mantel test. For each pair of individuals, the most recent divergence across the 100 phylogenetic trees was considered. The pairwise geographic distance along the Earth surface was computed from GPS coordinates, using the earth.dist function in the R package fossil (Vavrek 2011). For the Mantel test, the observed Spearman correlation coefficient was compared to those obtained with 9,999 permutated matrices. The R^2^ was extracted from a linear model.

### Inferring the rates of gains and losses of LGT

To estimate the times of origin, each LGT was recoded as a presence/absence character and mapped onto a time-calibrated phylogenetic tree. These analyses were performed on a per gene basis and not at the genomic block level because genes can be independently lost and blocks in less contiguous genomes will be more fragmented. To allow each LGT to evolve along the phylogenetic tree that best explained its history, the likelihood was estimated using the ace function with an asymmetrical substitution matrix (ARD model) in the ape package (Paradis and Schliep 2019) for all 100 phylogenetic trees, and the tree producing the highest likelihood was selected. Stochastic mapping was then performed to map the history of the LGT using the make.simmap function in the phytools package (Revell 2012). Independently for each of the five reference genomes, the time of acquisition along branches connecting the root to the tip corresponding to the reference genome was recorded for each of the LGT detected in the reference genome. If more than one acquisition was inferred along these branches, the history of the LGT was considered as ambiguous and the LGT was not included in the rate analyses. The process was repeated 100 times, producing 100 sets of time of origin for each of the LGT in each of the five reference genomes. These repeats represent pseudoreplicates. Note that, for a given LGT, the number of acquisitions can vary among repeats of the stochastic mapping, so that the LGT is considered in some but not all repeats.

The number of LGT per million-year time slices was extracted from the distribution of times of origin, up to 11 million years ago. For each time slice, the number of acquired LGT averaged across the 100 replicates was used to estimate the rates of gains (*G*) and subsequent losses (*L*). For this purpose, an exponential decay equation was fitted using the nls function in R:

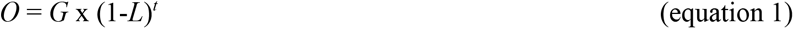

*O* is the number of LGT acquired in each time slice that were retained until the present and detected in the reference genome, and *t* is the average age of the time window. The rates *G* and *L* were estimated using the same approach independently for each of the 100 pseudoreplicates of stochastic mapping, and the 0.025 and 0.975 quantiles of their distributions were used to compute 95% confidence intervals.

## Analyses of native genes

From the initial 37-taxa trees, we extracted gene trees that were unlikely to have undergone LGT in their recent history. To be considered as such, a gene had to be present in at least *A. cimicina* and one of *A. angusta* and *A. semialata*, the *Alloteropsis* sequences had to be monophyletic, and they had to be placed in the phylogenetic tree within Paniceae (the tribe that contains *Alloteropsis*), but outside of the subtribes that do not contain *Alloteropsis*. Using this method, we identified 6,657 genes that existed in the common ancestor of *Alloteropsis* and have then been transmitted to some of its descendants. The presence of each of these genes in the five reference genomes was established based on read analyses, as described above for LGT. We identified 227 out of these 6,657 that were absent from at least one of the five reference genomes, and the presence/absence of each of these 227 genes across the 45 *Alloteropsis* individuals was established using read analyses, as described for the LGT.

The rate of losses of these native genes was inferred by mapping their origins on the phylogenetic trees, as described for the LGT. For each gene missing from a reference genome, its loss in the lineage leading to this genome was recorded. As before, if multiple losses were inferred along the branches leading to the reference genome, the gene was not considered when calculating rates. The number of losses across the 100 mapping replicates was then computed per 1 Ma time window. Because losses are not expected to be frequently recovered, the number of observed losses is not expected to decrease with their age, and if the rate of losses is constant, the number of losses per time window should not vary. We consequently computed the rate of losses as the mean number of observed losses across the eight most recent time windows, which represent the time during which *A. angusta* and *A. semialata* evolved separately.

## Data availability

raw sequence data, genome assemblies and annotations generated as part of this study are available from NCBI Genbank under BioProject PRJNA824797. All phylogenies and alignments have been made available on dryad: https://doi.org/10.5061/dryad.sbcc2fr9s. The Scripts used in this study are available from GitHub:

https://github.com/Sheffield-Plant-Evolutionary-Genomics/panLGT-Alloteropsis-2022.

## Author contributions

PR, CP, PAC, and LTD designed the study. PR, MEB, and LTD generated the genome data, MEB and LTD assembled and annotated the genomes, PR, LP, PAC, and LTD analysed the data. All authors interpreted the results and helped write the manuscript.

## Acknowledgements

This work was funded by the Natural Environment Research Council grant NE/V000012/1, PAC is funded by a Royal Society University Research Fellowship (grant URF\R\180022), and LTD is funded by a NERC fellowship (grant NE/T011025/1).

