## Supplementary Material for "Variation in the rate lateral gene transfers accumulate in a grass lineage"

#### Supplementary Methods

##### *Genome sequencing, assembly and annotation*

Illumina reads were filtered using NGSQCToolkit v.2.3.3 (Patel and Jain 2012) to remove adapter contamination and low quality reads (i.e. < 80% of sites with Phred > 20), and trimmed from the 3' end to remove low quality bases (Phred < 20). Read duplicates were removed using PrinSeq-lite v.0.20.3 (Schmieder and Edwards 2011). Finally, the reads were error-corrected using SOAPec v.2.01 (Luo et al. 2012). Pacbio reads were error-corrected with Proovread v.2.14.0 (Hackl et al. 2014) using the filtered Illumina reads. The amount of cleaned data used for assembling each reference genome is listed in Table S1.

Draft genomes were assembled for the three samples with long-read data using a hybrid strategy incorporating Illumina and Pacbio data. First, the filtered Illumina reads were assembled into contigs using SOAPdenovo2 v.2.04 (Luo et al. 2012) with the parameters *k-mer* = 65, *KmerFreqCutoff* = 10, *mergeLevel* = 3 (max), and *arcWeight* = 5. A hybrid assembly was then performed for samples with PacBio data using DBG2OLC v.11062016 (Ye et al. 2016) with the SOAPdenovo contigs and the error-corrected, trimmed Pacbio reads. RSA5-3 was assembled using the same parameters as AUS1 (Dunning et al. 2019)(MinOverlap = 60*;* KmerCovTh = 3; AdaptiveTh = 0.03), whereas parameters were modified to optimise N50 for ZAM1505-10 (MinOverlap = 15*;* KmerCovTh = 2; AdaptiveTh = 0.03) and TAN1-04B (MinOverlap = 15*;* KmerCovTh = 2; AdaptiveTh = 0.01). Scaffolding was then further improved using SSPACE v.3.0 (Boetzer et al. 2011) with the paired-end Illumina reads, followed by SSPACE-LongRead v.1.1 (Boetzer and Pirovano 2014) with the untrimmed Pacbio reads. As no long-read data was available for AANG_UGA4, the mate-pair libraries were used to generate scaffolds with SOAPdenovo2 as opposed to working with the contigs. Scaffolds of AANG_UGA4 longer than 500 bp were then arranged using Chromosomer v.0.1.3 (Tamazian et al. 2016) with default parameters and the chromosomes from the AUS1 genome as a reference. The completeness of each genome was assessed by checking for the presence of 4,896 Poales benchmarking universal single copy orthologs using BUSCO v.3.1.0 (Simão et al. 2015).

Genome annotation used the same *A. semialata* transcriptome database assembled to annotate the AUS1 genome (Dunning et al. 2019), that was constructed from nine published *A. semialata* transcriptomes (Dunning et al. 2017a). The transcriptome database was aligned to each reference genome assembly using PASA v.2.0.2 (Haas et al. 2003). For AANG_UGA4 the same approach was used, but with two previously published *A. angusta* transcriptomes (Dunning et al. 2017a). Annotation of the reference genomes was then performed using MAKER v.2.31.8 (Holt and Yandell 2011) with default parameters. This pipeline utilised RepeatMasker with a Poaceae repeat database (v.20160829), the above *Alloteropsis* reference transcriptome databases, and *Sorghum bicolor* (Sorbi1 [Paterson et al. 2009]) and *Setaria italica* (JGIv2.0 [Bennetzen et al. 2012]) protein datasets. For the *ab initio* gene prediction, both SNAP v.2013-02-16 [Korf et al. 2004] and Augustus v.3.2.3 (Stanke and Waack 2003) were used. The training files for SNAP and Augustus generated by Dunning et al. (2019) were used to run the analysis.

***Identification of laterally acquired genes***

To identify LGTs in each of the reference genomes, we used the phylogenetic method developed in Dunning et al. (2019) and Hibdige et al. (2021), with a few modifications. This approach uses a rapid phylogenetic screening method with a modest number of taxa (≤ 37 sp.), and builds a maximum likelihood phylogenetic tree with phyml v.21031022 (Guindon and Gascuel 2003) for each gene using only the top-hit Blastn match (based on the highest bit-score) from each species, with a minimum alignment length of 300 bp (not necessarily a single continuous Blast match). To be considered as a candidate LGT, the *Alloteropsis* sequence has to be nested within a distantly related group of grasses that is resolved in a coalescence species tree analysis (Dunning et al. 2019). A gene was classed as nested when its sister group, and also their combined sister group, belonged to the same group of grasses, and either of these groupings was supported by at least 50% of bootstrap pseudoreplicates. In addition, *Alloteropsis* genes were considered as candidate LGT if they were sister to a clade represented by a single species in our sampling (Aristidoideae, Danthonioideae, Chasmanthieae, Melinidinae and Tristachyideae), provided that their combined sister was as expected based on the species tree. After passing this initial filter, candidate LGTs were subjected to a more detailed phylogenetic analysis with an increased number of taxa (≤ 135 species), a more stringent bootstrap threshold (nesting supported by ≥ 70% of bootstrap replicates), and additional AU-topology tests. The full list of data and potential donor clades considered as part of this study can be found in Table S7. In contrast to Hibdige et al. (2021), the short-read datasets were added to the detailed phylogenetic analysis prior to the topology tests. This enabled us to maximise the number of donor clades considered, particularly for those lacking multiple transcriptome/genome datasets. We also used Blastn to identify and extract native and LGT sequences from the five *Alloteropsis* reference genomes, if they were present but not annotated, with sequences rescued from the *Alloteropsis* genome required to be at least 200 bp long. Finally, secondary LGT candidates were identified as in Dunning et al. (2019). Secondary LGT candidates are those that were not identified in our stringent phylogenetic pipeline, but are in close physical proximity in the *Alloteropsis* genomes to genes identified as LGT in our filtering and still produce gene trees that support the same LGT scenario and were potentially acquired as part of the same DNA fragment.

### Supplementary Results

#### Detection of LGT from five reference genomes

The primary LGT candidates (*sensu* Dunning et al. 2020) were identified using a stringent phylogenetic pipelines, which detected 138 protein-coding LGT across the five genomes (17 – 53 per genome; Table S3). We then analysed the genomic regions flanking these primary candidates and retrieved a total of 85 secondary LGT secondary (5 – 40 per genome; Table S3), which were related to the same group of donors as the adjacent primary candidates but were discarded during the stringent filtering. Finally, the other *Alloteropsis* genomes were scanned for genes homologous to either primary or secondary LGT candidates, and we retrieved 78 sequences corresponding to LGT that had been filtered out by the stringent pipeline and 17 that were unannotated in one genome (Table S3). 41 of the 64 LGT previously detected in AUS1 across two studies (Dunning et al. 2019; Hibdige et al. 2021) were re-identified here, with the 23 others being lost mainly during the topology tests or because secondary candidates identified purely based on read mapping were not considered here (Table S8). It is worth noting that the 14 lost due to the topology tests includes both the primary LGT which directly failed the test (n = 6), and the secondary candidates which were indirectly lost as a consequence (n = 8). Another 20 LGT were identified here in other genomes and then recovered in the genome of AUS1, and these were missed in previous studies (Table S8). In total, we identified 318 LGT, ranging from 34 in the genome of RSA5-3 to 130 in the genome of ZAM1505-10 (Table S3). These were combined into a non-degenerate list of 177 LGTs in *Alloteropsis* (Table S9), by first collapsing recent duplicates in the genome (removed 45 LGT; Table S3), and by only counting each LGT once even if it was present in multiple genomes.

We then assessed the presence of each one of the 177 LGTs within each of the five genomes based on the presence of corresponding reads, which can identify unannotated or unassembled genes. For 11 LGT, reads were consistently grouped with sequences other than the reference *Alloteropsis* LGT sequence, and these LGT were not considered further (Table S9). We thus assessed the distribution of the 166 remaining LGT among the five reference genomes. In 11 cases, reads from one genome corresponding to a LGT identified in another genome were detected (Table S9), leading to between 32 and 100 read-supported LGT per genome, with the highest number observed in the genome of ZAM1505 (Table S3). Only two of the 166 LGT were shared by all five accessions, 12 by four accessions, 15 by three accessions, 26 by two accessions, and 113 were found in only one of the five accessions. Of these, 20 were unique to AANG_UGA4, 11 to RSA5-3, 6 to TAN1-04B, 54 to ZAM1505-10, and 22 to AUS1 (Table S3).

##### *Multiple donors passed fragments of various sizes*

The 166 LGT were assigned to donor lineages based on their phylogenetic position, and transfers were detected from four of the nine core grass subfamilies (92.2% Panicoideae, 4.8% Chloridoideae, and 3.0% Danthonioideae; Table S4). The Panicoideae is the second-largest subfamily and includes over 3,500 species. Owing to the prevalence of data for this group, the donors from this subfamily can be further divided into sublineages, including Cenchrinae (n = 88 LGT), Andropogoneae (n = 54), Melinidinae (n = 7), Tristachyideae (n = 2), Panicinae (n = 1), and the *Lasiacis*-*Cyrtococcum* group (n = 1; Table S4). Overall, the taxonomic distribution largely agrees with previous patterns, with a higher number of LGT from closer relatives (Dunning et al. 2019; Hibdige et al. 2021). The identity of the donor slightly varies among the reference genomes, with LGT received from Danthonioideae restricted to the South African RSA5-3 (Table S4).

Distinct protein-coding LGT were assigned to the same genome block if they were adjacent in any of the five reference genomes (Table S9 & 10). In total, the 166 LGT could be assigned to 82 different genomic blocks, with the largest containing 12 distinct LGT in the ZAM1505-10 genome (block 68; Table S4). A total of 45 LGT appeared as singletons in the assembled genomes (Table S4). All genes from the same fragment were assigned to the same donor, with one exception (Figure S1; Table S4). Block 63 contains three LGT, and the first two are assigned to Cenchrinae while the last one is assigned to Andropogoneae and is present in more *Alloteropsis* accessions (Figure S1). The third gene is present in three distinct contigs in the assembled genome of TAN1-04B that likely represent post-transfer duplicates, and only one of them is joined with the two other LGT (Table S4 & S9). These patterns might result either from a misassembly or from distinct LGT clustering in the genome after independent transfers, as previously suggested in *Hordeum* (Mahelka et al. 2021). The presence of each of the 166 LGT was then assessed for 40 other accessions belonging to the three *Alloteropsis* species. The two LGT identified in all five reference genomes were identified in 44 out of 45 accessions, and the number of accessions bearing each of the other LGT ranged from 40 to 1, with 5 LGT identified only in one individual (Table S4).

**Phylogenetic distribution of LGT suggests few old LGT and many recent ones**

To capture both analytical uncertainty and different gene histories, we inferred 100 dated phylogenies from random sets of five single-copy orthologs. The inferred species tree varied slightly among the different gene sets, although *A. semialata* and *A. angusta* were always monophyletic (Figure 1). Within *A. angusta*, we recovered the two main clades corresponding to erect and decumbent ecotypes (Curran et al. 2022), with some gene discordance compatible with gene flow after their split (Figure 1). Similarly, the four nuclear clades previously described within *A. semialata* (clades I-IV) (Olofsson et al. 2016; Bianconi et al. 2020) were recovered, with gene discordance mainly concerning the relationships among these clades (Figure 1). In particular, clade

II was placed with similar frequency as sister to clade I and as sister to clades III+IV, reflecting hybridization at its base (Bianconi et al. 2020).

The distribution of LGT largely mirrored the phylogenetic relationships. Three genes are present in all *A. cimicina* and most *A. semialata* and *A. angusta* accessions (Figure 1), and long branches leading to the LGT of different accessions are compatible with an ancient acquisition followed by the accumulation of novel mutations in each species (Figure 2). These three genes were acquired from Chloridoideae grasses, and two are in the same genomic block (Figure S1; Table S4). A few of the other LGT were shared among most *A. angusta* and *A. semialata* accessions, with more specific to each species or subclade of *A. semialata* (Figure 1). Of the LGT specific to *A. angusta*, most were widely shared within the species, with one specific to AANG_UGA4 and one shared with only one other accession (Figures 2 and S1). Several of the LGT detected in AUS1 were restricted to other accessions from Australia and the Philippines, as previously reported (Dunning et al. 2019; Olofsson et al. 2019). None of the LGT detected in RSA5-3 or TAN1-04B were specific to these accessions, being widely shared among other members of their clades (Figure 2). All the LGT found in TAN1-04B were also found in the other individual from the same population with sequence data available (TAN1-04A; Table S4). The pattern differs for ZAM1505-10, which possesses three LGT found in no other individual, and 19 shared only with the other Zambian individuals from clade III (Figures 2 and S1; Table S4).

Overall, the phylogenetic patterns show that few LGT are shared by most accessions and thus likely ancient, while most are specific to subgroups, and likely happened recently. Comparisons among the five reference genomes suggests more recent LGT in ZAM1505, which is compatible with the highest number of LGT unique to this accession.

**Further investigation indicates some genes have been independently acquired in different *Alloteropsis* lineages**

Most of the genes within each genomic block have comparable distributions among accessions (Figure S1), although we identify a few notable exceptions. One of the largest fragments includes twelve protein coding genes (block 68), 11 of which are restricted to Zambian and Tanzanian accessions, while the other is shared with Australian accessions (LGT-037; Figure S1). In the original phylogeny of LGT-037, the genes from the reference genomes ZAM1505-10 and AUS1 were grouped and nested within Andropogoneae, but with important divergence between them. We added sequences from a variety of *Themeda* accessions retrieved from other papers (Dunning et al. 2017b; Arthan et al. 2021; Dunning et al. 2022), a genus of Andropogoneae that was previously identified as the donor of LGT detected in AUS1 (Dunning et al. 2019). In the denser phylogenetic tree, the gene from ZAM1505-10 was nested within *Themeda triandra*, while the gene from AUS1 was placed outside of *Themeda* and closely related genera (Figure S6). This phylogenetic pattern suggests that homologous genes were transferred from distinct Andropogoneae species independently to Zambian and Australian *A. semialata*. As a result, we placed the Australian gene into its own block (block 80). We similarly added sequences from *Themeda* and close relatives to the phylogeny of the original LGT-42 (later split between block 16 and the first gene of block 69), which was also distributed more widely than the other genes within block 69 (Figure S1). As for LGT-037, the sequence from ZAM1505-10 was nested within *T. triandra*, while the sequence from MRL48 was placed outside of *Themeda* (Figure S7), pointing to two different LGT of homologous genes. Because of the shortness of the fragment, the relationships however received low bootstrap support (Figure S7). We reanalyzed the reads corresponding to these two LGT, assigning them to one of the LGT resulting from independent transfers, which were considered as independent LGT in all other analyses (LGT-037a versus LGT-037b and LGT-042a versus LGT-042b). This led to a total of 168 LGT distributed among 84 genomic blocks, 47 of which are singletons.

**Introgression likely spread recent LGT among accessions**

Some of the LGT have a patchy distribution among the phylogenetic groups (Figure 1), which could result from recurrent losses after a relatively ancient acquisition, or introgression after a recent acquisition, as previously shown for some LGT (Olofsson et al. 2016; Dunning et al. 2017a). The second scenario is supported in several cases by the sharing of LGT among geographically close accessions belonging to distinct lineages. For example, the ZAM1720 *A. angusta* from the North of Zambia shares six LGT (representing two genomic blocks; blocks 48 and 49 in Figure S1) with clade II individuals from the South of Tanzania (closest one collected 206km away), and these LGT are absent from other *A. angusta* accessions (Figures 2 and S1). Similarly, the individual ZAM1507 from Zambia shares LGT, otherwise absent from clade II, with clade III individuals from Zambia and Tanzania (Figure 2), including the 12-LGT genomic block 68 (Figure S1). This individual comes from a mixed population of clade II and clade III individuals, for which admixture was evidenced (Olofsson et al. 2021). Finally, several LGT blocks are shared among different African clades, but are absent from Asia and Australia (e.g. blocks 21, 22, 50, 52, 53, 55, 63, 65), which could result from losses during the migration to Australia or introgression within Africa after the split of the Asian lineage. The phylogenetic signal retracing the sequential acquisition of LGT is therefore partially blurred by likely introgression following the original transfers.

**LGT are lost faster than native genes**

The rate of gene loss was estimated based on 6,657 native genes present in *A. cimicina* and at least one of the five reference genomes. Of these 6,657 genes, 227 were absent from at least one of the five genomes. The distribution of these 227 genes among the 45 accessions was used to estimate, for the lineages leading to each of the five reference genomes, the number of losses per Ma, as done for the LGT. Many of these gene losses have a patchy distribution among accessions, likely reflecting recurrent losses complicated by the fact that gene absence is a recessive character, so that heterozygous individuals cannot be detected with our approach. As a consequence, the origin of the loss could not be estimated for many missing genes (86/170 in AANG_UGA4, 24/35 in RSA5-3, 26/34 in TAN1-04B, 26/39 in ZAM1505-10, and 27/38 in AUS1). The distribution of the losses for which ages could be inferred was mostly flat through time, although the high rate of losses observed in AANG_UGA4 decreased in the last 2 Ma (Figure S4). The means over the last 8 Ma (time since the split of *A. angusta* and *A. semialata*; Figure 1) ranged from 0.7 gene per Ma in TAN1-04B to 8.4 in AANG_UGA4 (Figure S4). Assuming that the genes whose losses could not be inferred are evenly distributed through time, these numbers would scale to between 3.0 genes per Ma in TAN1-04B and 16.9 in AANG_UGA. Given that we analysed 6,657 genes, these numbers indicate that between 0.04 and 0.06 % of native genes were lost per Ma in *A. semialata*, while 0.25 % were lost in *A. angusta*. The increase of gene loss in *A. angusta* is not associated with the reduced completeness of the genome assembly as this analysis was performed on the unassemble short-read data.

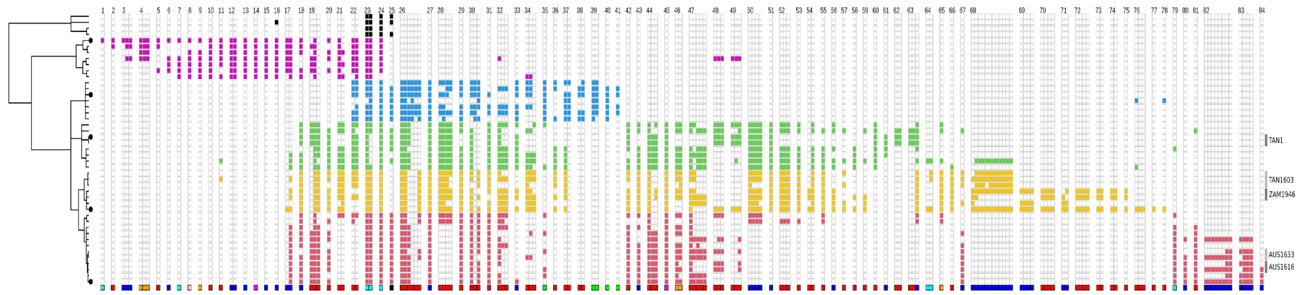


**Fig. S1. Distribution of laterally acquired genomic blocks.** The distribution of 168 genes resulting from lateral gene transfers (LGT) is indicated, with presence shown by squares coloured by clade; black = *A. cimicina*, purple = *A. angusta*, blue = clade I of *A. semialata*, green = clade II of *A. semialata*, orange = clade III of *A. semialata*, red = clade IV of *A. semialata*. Empty columns separate genomic blocks, which are numbered at the top. The identity of the donor is shown at the bottom of each column; blue = Andropogoneae, red= Cenchrinae, orange = Melinidinae, cyan = Chloridoideae, magenta = Tristachyideae, green = Danthonioideae, black = Cyrtococcum-Lasiacis group, pink = Panicinae. A consensus of the 100 phylogenies shown in Figure 1 is presented on the left, with black circles at tips highlighting accessions for which a complete genome is available. Populations with multiple individuals are delimited and named on the right. See Table S4 for details

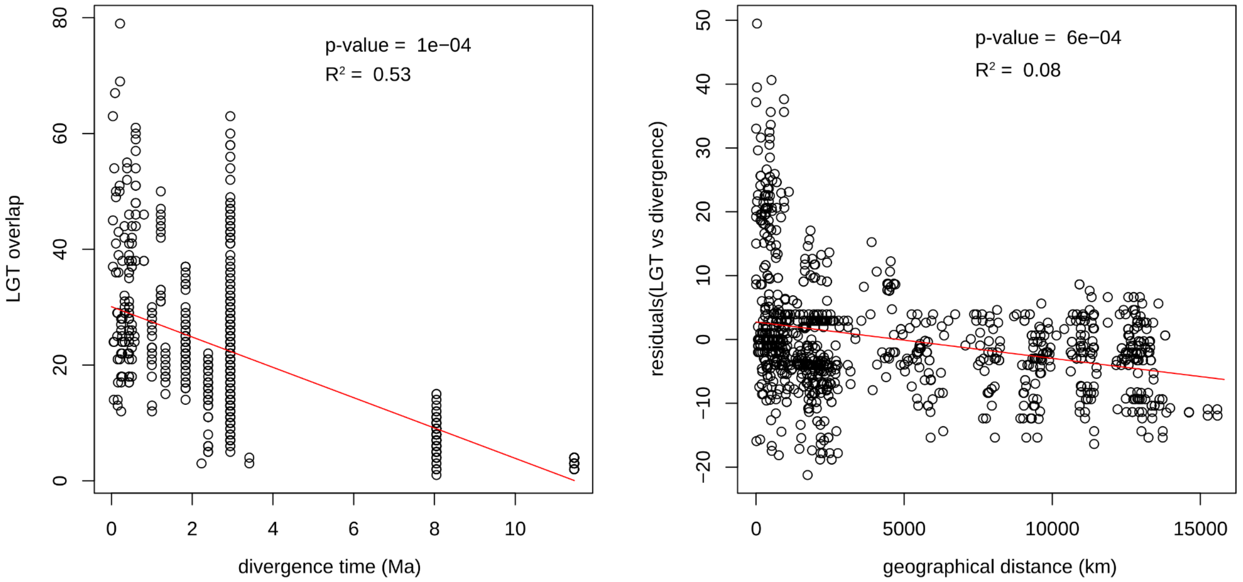


**Fig. S2. Effects of history and geography on the distribution of lateral gene transfers among accessions.** For each pair of individuals, the number of shared lateral gene transfers (LGT) is plotted as a function of the divergence time (in million years, Ma; left panel). The residuals of the regression between these two variables are then plotted as a function of the pairwise geographical distances (in km along the Earth's surface; right panel). The p-values, based on Mantel tests, are shown on the graphs, together with the $R^2$ calculated using a linear regression. The regression lines are represented in red.

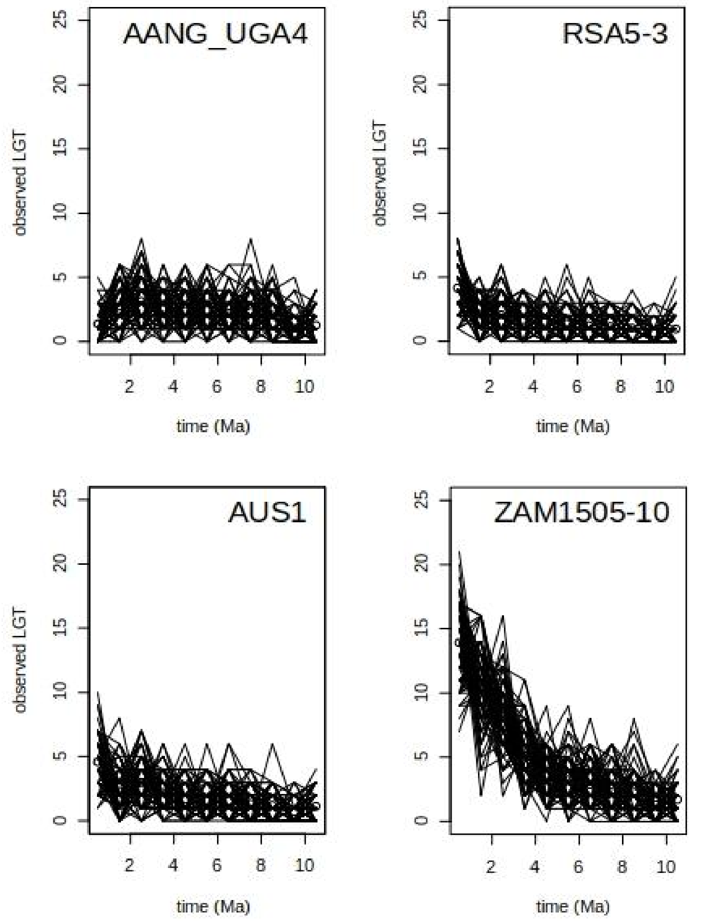


**Fig. S3: Patterns of gains of laterally acquired genes through time for pseudoreplicates.** For each reference genome, the number of genes resulting from lateral gene transfers (LGT) per 1 Ma is indicated for 100 pseudoreplicates, with values from the same pseudoreplicate connected with lines.

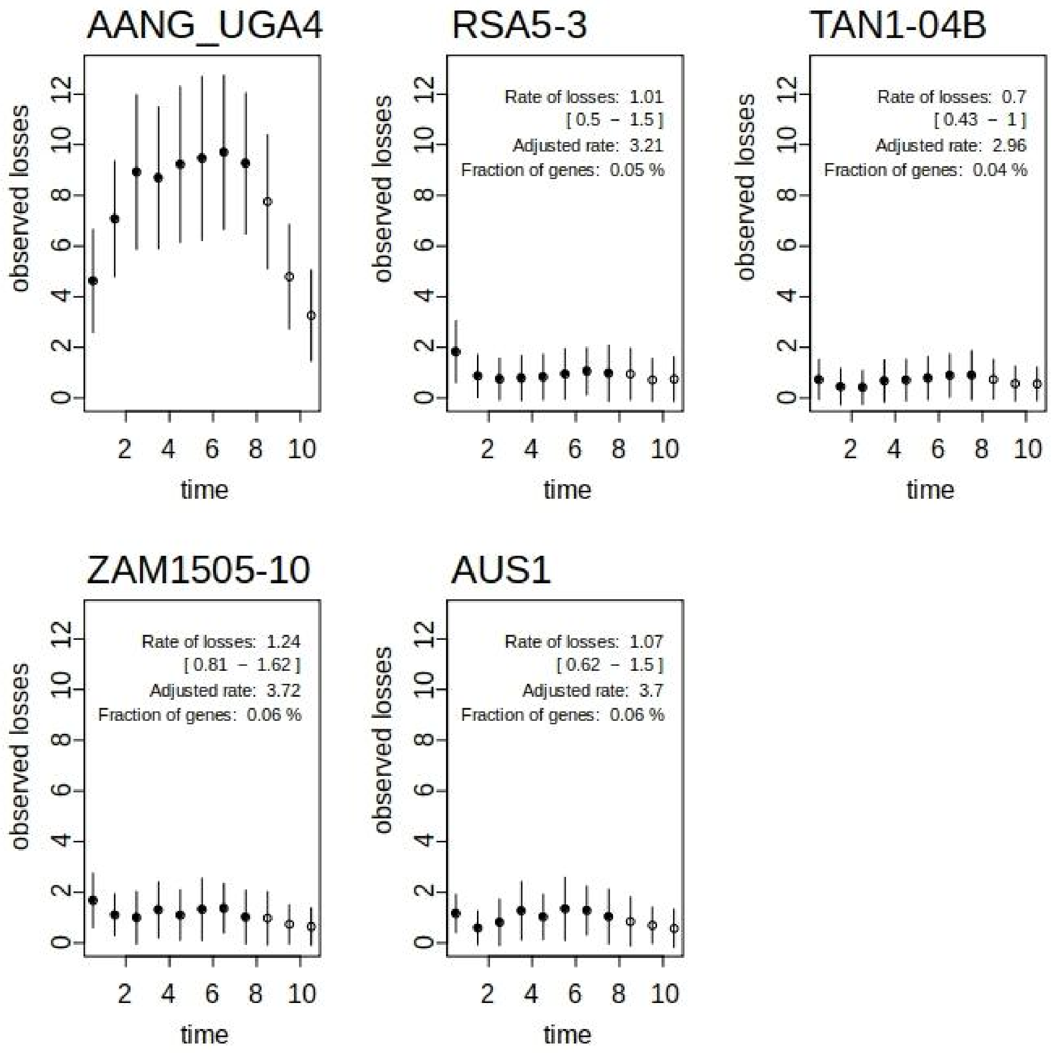


**Figure S4: Losses of native genes through time.** For each of the five reference genomes, the mean number of losses of native genes inferred for each 1 Ma time window is represented with dots. The error bars show the 95% interval among 100 replicates. Points used to calculate the mean over the last 8 Ma are in black. For all *A. semialata* genomes, the inferred rate of losses is indicated, with the 95% confidence interval based on 100 replicates in square brackets. The rate adjusted for missing data is also shown, together with the fraction of the 6,657 genes represented by the adjusted rate. For AANG_UGA4, the rate of losses was 8.37 genes per Ma [7.38-9.63] and the adjusted rate was 16.93 genes per Ma, which represents 0.25% of the 6,657 genes considered.

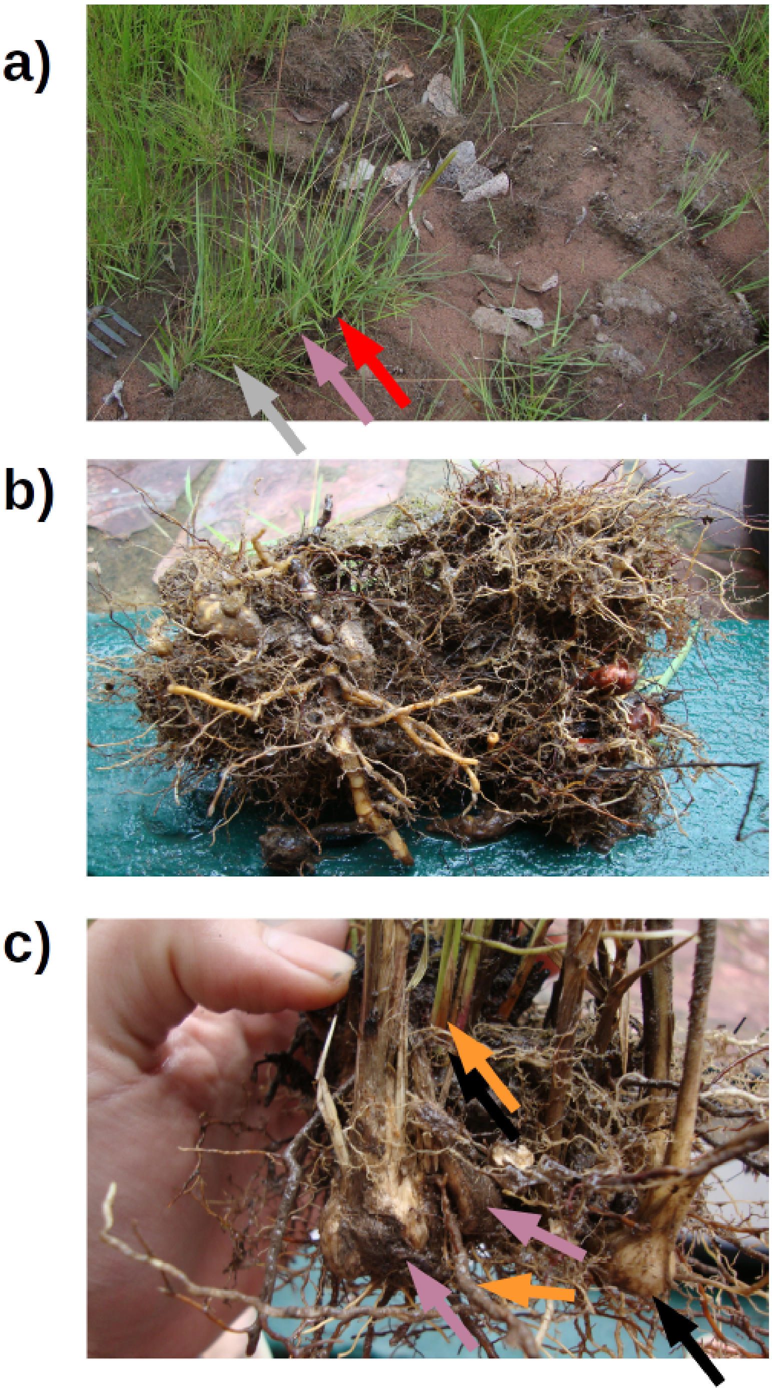


**Fig. S5: Examples of multispecies clumps in Zambia.** a) Picture from population ZAM1946 (same location as ZAM1710 in [Olofsson et al. 2021]). *Alloteropsis semialata* (purple arrow) grew together with *Setaria* sp. (Cenchrinae, red arrow) and an unidentified grass (grey arrow). b) Multispecies clump seen from below. The individual ZAM1946-12 of *A. semialata* grew with its roots and bulbs entangled with those of numerous other species. c) Multispecies clumps seen after removing the soil. Individual ZAM1946-10 of *A. semialata* (purple arrow) grew entangled with other species, including *Brachiaria* sp. (Melinidinae, orange arrow) and other unidentified species (black arrow).

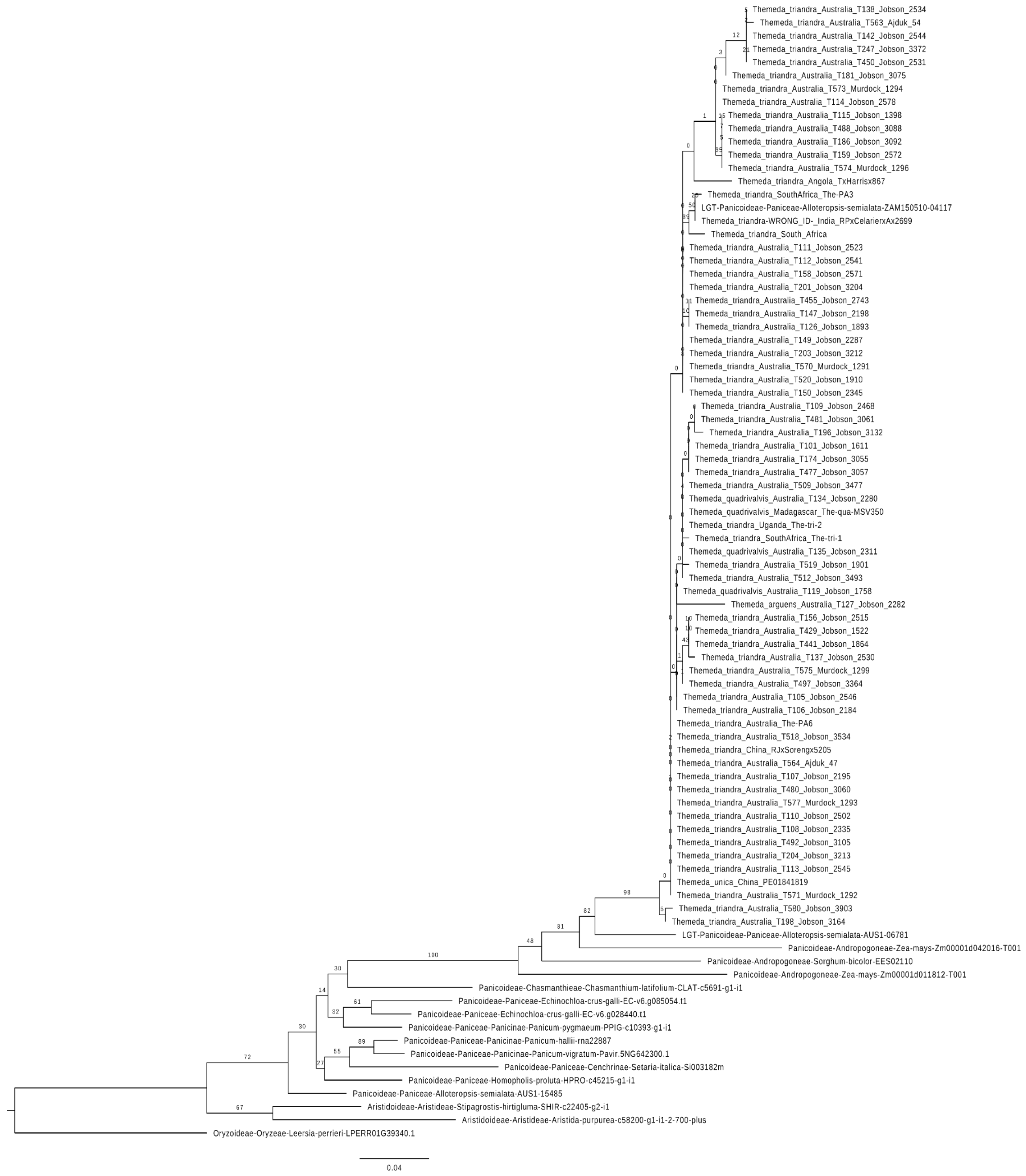


**Fig. S6: Phylogenetic tree of gene ZAM1505-10-04117 and homologs.** The gene is the first from block 68 (LGT-037; Table S4). A maximum-likelihood phylogenetic tree was inferred after incorporating a dense sampling of samples from *Themeda* and relatives. Bootstrap support values are indicated near nodes.

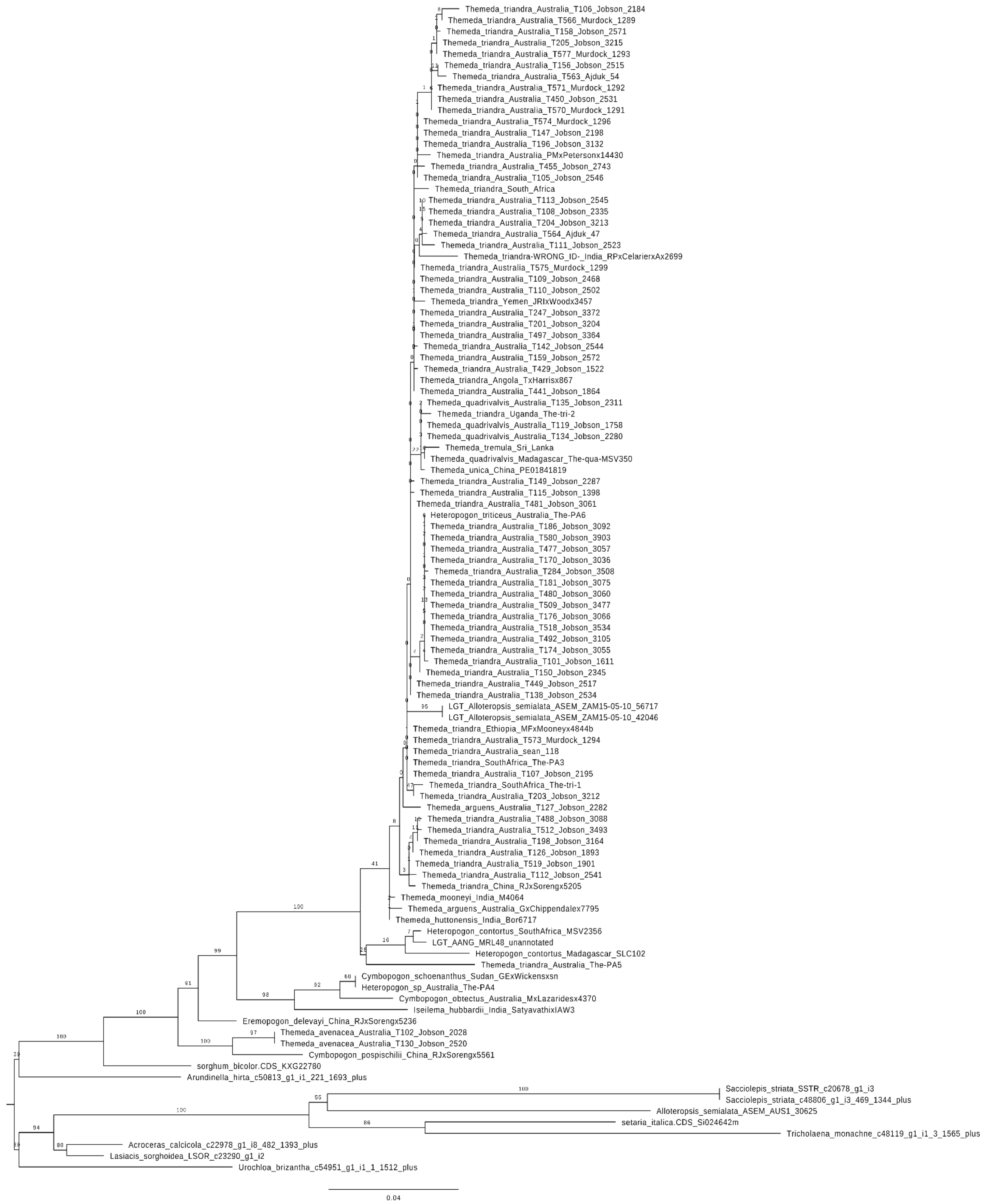


**Fig. S7: Phylogenetic tree of gene ZAM1505-10-42046 and homologs.** The gene is the first from block 69 (LGT-042; Table S4). A maximum-likelihood phylogenetic tree was inferred after incorporating a dense sampling of samples from *Themeda* and relatives. Bootstrap support values are indicated near nodes.

**Table S1:** List of cleaned sequence datasets used for genome analyses

| Library | AANG UGA4 | RSA5-3 | TAN1-04B | ZAM1505-10 | AUS1 |
|---|---|---|---|---|---|
| Illumina Paired-end [350bp insert – 125bp reads] | 41.2 Gb | 40.8 Gb | - | - | 51.8 Gb |
| Illumina Paired-end [550bp insert – 125bp reads] | - | - | - | - | 46.2 Gb |
| Illumina Paired-end [550bp insert – 250bp reads] | 43.6 Gb | 47.9 Gb | 49.5 Gb | 49.9 Gb | 64.0 Gb |
| Illumina Mate-pair [5kb insert – 125 bp reads] | 9.8 Gb | 8.7 Gb | - | - | 9.9 Gb |
| Illumina Mate-pair [10kb insert – 125 bp reads] | - | 18.8 Gb | - | - | 9.6 Gb |
| Illumina Mate-pair [10kb insert – 250 bp reads] | 17.6 Gb | - | - | - | - |
| Pacbio | - | 6.3 Gb | 11.7 Gb | 9.3 Gb | 6.2 Gb |
| PacBio N50 | - | 5.2 kb | 4.9 kb | 4.3 kb | 5.3 kb |

**Table S2:** Genome assembly statistics for the accessions of *Alloteropsis* used in this study.

| | *A. angusta* | *A. semialata* | | | |
|---|---|---|---|---|---|
| **Metric** | **AANG_UGA4** | **RSA5-3** | **TAN1-04B** | **ZAM1505-10** | **AUS1** |
| Estimated genome size (1C/Gb)[1] | 0.97 | 0.87 | 1.01 | 1.09 | 1.10 |
| Total assembly length (Gbp) | 1.54 [0.93][2] | 0.62 | 0.75 | 0.87 | 0.75 |
| Relative assembly size | 158.76% [95.88%][2] | 71.26% | 74.25% | 79.81% | 68.18% |
| Proportion of 'N' | 65.49% [40.28%][2] | 5.85% | 2.68% | 4.22% | 5.82% |
| Scaffold N50 (Mb) | 106.76 [0.02][2] | 0.18 | 0.17 | 0.07 | 82.05 |
| Longest scaffold (Mb) | 161.46 [0.13][2] | 0.99 | 1.07 | 0.60 | 99.96 |
| Number of scaffolds | 85,825 [179,640][2] | 5,965 | 7,979 | 19,813 | 686 |
| Annotated genes | 46,005 | 47,028 | 51,145 | 67,506 | 45,144 |
| BUSCO analysis assembly (n = 4,896 genes): | | | | | |
| Complete | 77.1% | 86.7% | 90.5% | 91.9% | 88.1% |
| Complete single | 65.9% | 78.3% | 82.1% | 66.4% | 82.9% |
| Complete duplicated | 11.2% | 8.4% | 8.4% | 25.5% | 5.2% |
| Fragmented | 4.6% | 2.1% | 1.7% | 1.9% | 1.7% |
| Missing | 18.3% | 11.2% | 7.8% | 6.2% | 10.2% |
| BUSCO analysis coding sequences (n = 4,896 genes): | | | | | |
| Complete | 67.4% | 82.0% | 85.3% | 87.8% | 80.9% |
| Complete single | 54.8% | 73.4% | 76.9% | 61.5% | 75.3% |
| Complete duplicated | 12.6% | 8.6% | 8.4% | 26.3% | 5.6% |
| Fragmented | 9.3% | 3.9% | 3.3% | 3.5% | 4.6% |
| Missing | 23.3% | 14.1% | 11.4% | 8.7% | 14.5% |

[1] Extracted from Bianconi et al. (2020).

[2] Value prior to being scaffolded using the AUS1 genome as a reference

**Table S3: Summary of LGT numbers per genome**

| Individual | Primary | Secondary | Others | Total | Collapsed | Read supported | Unique | Age missing[a] |
|---|---|---|---|---|---|---|---|---|
| AANG_UGA4 | 19 | 6 | 11 | 36 | 35 | 32 | 20 | 8 (25%) |
| RSA5-3 | 17 | 5 | 12 | 34 | 32 | 33 | 11 | 14 (42%) |
| TAN1-04B | 24 | 14 | 19 | 57 | 49 | 45 | 6 | 19 (42%) |
| ZAM1505-10 | 53 | 40 | 37 | 130 | 100 | 100 | 54 | 44 (44%) |
| AUS1 | 25 | 20 | 16 | 61 | 57 | 50 | 22 | 28 (56%) |
| Total | 138 | 85 | 95 | 318 | 273 | 260 | 113 | 113 |

[a] Median across 100 replicates of the number of LGT for which an age of acquisition age could not be estimated.

**Tables S4 - S10 are available as individuals tabs in the TableS4-S10.xlsx file**

**Table S4:** Properties and distribution of detected LGT

**Table S5**: SwissProt LGT annotations

**Table S6**: Gene ontology (GO) enrichment analysis results

**Table S7**: Datasets used in tree construction

**Table S8**: Comparison of LGT detected from the AUS1 genome assembly in different studies

**Table S9**: Detected LGT and number of reads assigned to each of them

**Table S10**: Detailed information on primary LGT candidates
